# Genomic convergence of locus-based GWAS meta-analysis identifies DDX11 as a novel Systemic Lupus Erythematosus gene

**DOI:** 10.1101/2020.04.17.047332

**Authors:** Mohammad Saeed, Alejandro Ibáñez-Costa, Alejandra María Patiño-Trives, Eduardo Collantes Estevez, María Ángeles Aguirre, Chary López-Pedrera

## Abstract

Genome-wide association studies (GWAS) of systemic lupus erythematosus (SLE) explain only ∼15% of genetic risk, indicating genes of modest effect remain to be discovered. Association clustering methods such as OASIS are more apt at identifying modest genetic effects. 410 genes were mapped to previously identified OASIS GWAS SLE loci and investigated for expression in SLE GEO datasets. GSE50395 dataset from Cordoba was used for validation. Blood eQTL for significant SNPs in SLE loci and STRING for functional pathways of differentially expressed genes was used. Confirmatory qPCR on monocytes of 12 SLE patients and controls was performed. We identified 55 genes that were differentially expressed in at least 2 SLE GEO datasets with all probes directionally aligned. *DDX11* was downregulated in both GEO (P=3.60E-02) and Cordoba (*P*=8.02E-03) datasets and confirmed by qPCR (*P*=0.001). The most significant SNP, rs3741869 (*P*=3.2E-05) in OASIS locus 12p11.21, containing *DDX11*, was a cis-eQTL regulating *DDX11* expression (P=8.62E-05). *DDX11* interacted with multiple genes including *STAT1/STAT4*. Genomic convergence with OASIS and multiple expression datasets identifies novel genes. *DDX11*, RNA helicase involved in genome stability, is repressed in SLE.

**Summary Statement:** More than 100 genes for SLE have been identified but they explain only ∼15% of heritability. GWAS are challenged by risk genes of modest effect. Using locus-based GWAS mapping and multiple gene expression replications, *DDX11* was identified as a novel SLE gene. *DDX11*, repressed in SLE, may be used as a clinical diagnostic tool.

## Introduction

Systemic lupus erythematosus (SLE) is a diverse complex disorder, manifesting more like syndromes than a singular disease (Saeed. 2017a). SLE could therefore be thought of as a mixture of multiple resembling phenotypes, each a result of a separate mutation, pooled in genome-wide association study (GWAS) cohorts (Saeed. 2017b). Finding a particular gene then depends on the enrichment of a causal mutation carrying haplotype in the study sample.

Genotyping a multitude of single-nucleotide polymorphisms (SNPs) in GWAS leads to the multiple testing problem, preventing the true association signal from being distinguished from random noise. Despite the identification of large number of genes in multiple comprehensive SLE GWAS and candidate gene studies, they together explain only 15% of SLE heritability (Bentham et al. 2015; Morris et al., 2016).

OASIS is a linkage disequilibrium (LD) clustering method that can be used to mine existing complex disease GWAS datasets for new genes (Saeed, 2017b, Saeed, 2018). OASIS provides an alternative to increasing sample size for GWAS by composite analysis unifying two aspects of the LD phenomenon - strength of association and number of surrounding significant SNPs. A genomic convergence approach (Liang et al., 2009) mapping genetic association signals on expression data and other biological studies can then be used for verification of particular genes.

Previously, OASIS analysis of two dbGAP GWAS datasets (6077 subjects; 0.75 million SNPs) identified thirty SLE associated loci (Saeed, 2017b). In the present study, 410 genes were mapped to these loci and investigated for gene expression in three SLE GEO datasets. A fourth expression dataset from Cordoba, Spain was used for validation. This study showed that *DDX11*, a RNA helicase involved in genome stability, is repressed in SLE. Thus, a genomic convergence approach with OASIS, and expression studies on multiple datasets can help identify novel SLE genes.

## Results

### Genomic Convergence

SNIPPER located 410 genes (Table S1) in 30 SLE loci identified by OASIS (Saeed 2017b) and these were tested in 3 GEO expression datasets. This search identified over 1000 expression probe sets in GEO datasets GSE30153, GSE13887 and more than 500 probe sets in GEO dataset GSE10325 for the reference gene list. Significant T-tests for these probe tests identified, 103 probes in dataset 4193 (GSE30153), 141 probes in dataset 4719 (GSE13887) and 66 probes in dataset 4185 (GSE10325). Additional 215 probes were found to be significant in GSE10325: T-cells, B-cells and Monocytes datasets. Hence, a total of 525 statistically significant probe sets in six (3+3) GEO datasets were finalized. This composite probe list was matched against the SNIPPER reference gene list of 410 genes, to remove those that the GEO query included, however, were not found in the OASIS SLE loci. Hence, 187 unique genes in the 6 GEO datasets were identified that had significant change in expression and were located in the 30 SLE loci (Table S2).

### Expression Validation

Of the 187 candidate genes, 55 genes were found to be differentially expressed in at least 2 GEO datasets with all probes directionally aligned (Table S3). Four Genes crossed the false discovery rate (FDR) viz *STAT1, IFIH1, NRF1* and *TTC9* (Table 1). *DDX11*, located in OASIS locus 19 on chromosome 12, was downregulated, however it did not cross the FDR for expression. In the Cordoba expression dataset, GSE50395, 518 genes were found to have differential expression. Of these 10 genes were found in the reference list of 410 genes in OASIS SLE loci (Table 2). *BARHL1* had the most significant expression change (*P*=8.9 × 10^−5^) and *DDX11* had the second highest expression change (*P*=8 × 10^−3^).

**Table 1.** Top SLE candidate genes by genetic association and gene expression validation. Of 55 genes that showed statistically significant and consistent expression change in at least 2 GEO datasets, 4 crossed the FDR. *STAT4* and *IFIH1* are known SLE genes, however *NRF1* and *TTC9* need further data for validation. *DDX11* did not cross the FDR.

| Genetic Association |  |  |  |  |  | Gene Expression |  |  |  |  |
| --- | --- | --- | --- | --- | --- | --- | --- | --- | --- | --- |
| *Gene* | Chr: Position (bp) | Locus | Quad (Db1, Db2) | Max_SNP (Db1, Db2) | Max_P (Db1, Db2) | Significant Probes | T_Ave (P) | T-Min (P) | FDR (P) | Ave_Fold Change |
| *STAT1* | 2: 191838203...192060779 | 3 | A, B | rs7574865, rs3771327 | 13.05, 4.06 | 26 | 4.64E-03 | 7.75E-08 | 9.26E-04 | 36.19 |
| *IFIH1* | 2: 161182794...163311201 | 2 | C, B | rs6755940, rs13023380 | 3.12, 4.16 | 8 | 1.41E-02 | 7.19E-05 | 9.43E-04 | 25.02 |
| *NRF1* | 7: 128393949...132065378 | 9 | B, B | rs4728142, rs10488631 | 6.14, 10.78 | 6 | 3.00E-02 | 7.98E-05 | 9.62E-04 | 0.67 |
| *TTC9* | 14: 69831036...73507392 | 25 | C, C | rs7141009, rs2052219 | 2.73, 3.09 | 3 | 1.93E-02 | 1.20E-04 | 9.80E-04 | 0.76 |
| *DDX11* | 12: 30383902...31288883 | 19 | C, B | rs10843682, rs3741869 | 1.89, 4.50 | 2 | 3.90E-02 | 3.60E-02 | 2.94E-03 | 0.7 |

**Table 2.** Genomic Convergence of 410 reference genes in Cordoba Expression Dataset. Ten of 518 genes with expression change in the Cordoba dataset, were present in the reference gene list (410 genes). *DDX11* had the second highest expression change (*P*=8 × 10^−3^).

| Gene | FC Abs | P-value | Chr | OASIS Locus | Locus Position | Max_SNP (db1, db2) | Max_-log(P) (db1, db2) | Quadrant |
| --- | --- | --- | --- | --- | --- | --- | --- | --- |
| *BARHL1* | -2.66 | 8.85E-05 | 9q34.13 | 13 | 9: 135083870..135453438 | rs11243676, rs1185995 | 5.58, 4.97 | B, B |
| *DDX11* | -2.08 | 8.02E-03 | 12p11.21 | 19 | 12: 30383902..31288883 | rs10843682, rs3741869 | 1.89, 4.50 | C, B |
| *TRIM72* | -2.30 | 1.79E-02 | 16p11.2 | 27 | 16: 31193941..31527819 | rs11574637, rs9888739 | 6.27, 4.00 | B, B |
| *RBMS1* | -2.88 | 2.39E-02 | 2q24.2 | 2 | 2: 161182794..163311201 | rs6755940, rs13023380 | 3.12, 4.16 | C, B |
| *CDKN1B* | 2.47 | 2.76E-02 | 12p13.1 | 18 | 12: 12221429..13759033 | rs7980903, rs4764011 | 4.39, 3.26 | B, C |
| *ADAM21* | -3.19 | 2.97E-02 | 14q24.2 | 25 | 14: 69831036..73507392 | rs7141009, rs2052219 | 2.73, 3.09 | C, C |
| *ZNF385D* | -4.66 | 3.89E-02 | 3p24.3 | 5 | 3: 21523341..23226297 | rs3860579, rs680930 | 3.85, 3.06 | B, C |
| *LOC100286922* | -2.25 | 4.01E-02 | 2q37.1 | 4 | 2: 234514762..234825092 | rs17863787, rs4663968 | 3.17, 2.61 | C, C |
| *LOH12CR2* | 2.13 | 4.34E-02 | 12p13.2 | 18 | 12: 12221429..13759033 | rs7980903, rs4764011 | 4.39, 3.26 | B, C |
| *ELF5* | -2.21 | 4.47E-02 | 11p13 | 15 | 11: 34821947..35251317 | rs519858, rs2785197 | 2.57, 3.41 | C, C |

### eQTL analysis and qPCR

In 6 of 10 genes identified in the Cordoba dataset (GSE50395) that mapped onto OASIS loci, eQTLs were detected. The most significant SNP rs3741869 (GWAS P=3.2E-05) in OASIS locus 19 (chromosome 12p11.21), containing the gene *DDX11*, was found to be a *cis*-eQTL regulating the expression of *DDX11* (P=8.62E-05). The other GWAS significant SNPs modulated expression of genes not significant in GEO or Cordoba datasets (Table S4). Hence, *DDX11* was the only gene that could be functionally verified in this study by genomic convergence.

Expression of *DDX11* was confirmed with qPCR in monocytes isolated from 12 SLE patients and 12 healthy donors. As shown in Figure 2, *DDX11* was significantly downregulated in SLE patients (*P*=0.0010). We adjusted the data as logarithms, however despite any normalization technique the results remained the same.

**Figure 1.**
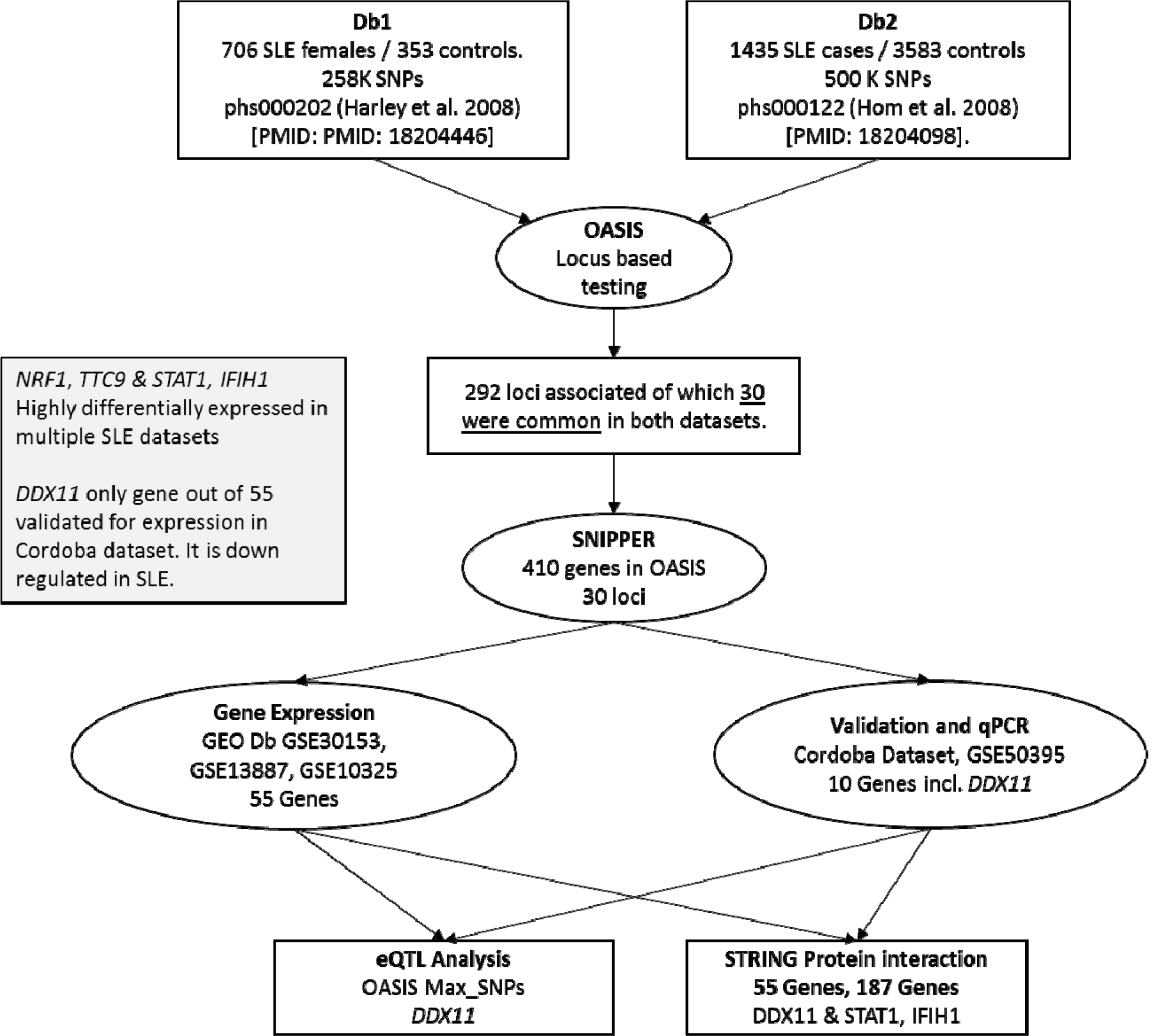
Methodology for gene discovery and validation. Genomic convergence methodology employed in this study. Two GWAS (Db1 and Db2) were analyzed using OASIS locus-based testing, which identified 30 overlapping SLE loci. SNIPPER was used to identify 410 genes in these 30 loci and this reference gene list was used for gene expression analysis in 3 SLE GEO datasets and 1 dataset from Cordoba, Spain. DDX11 was the only gene identified across all expression datasets with consistently significant expression change in SLE.

**Figure 2.**
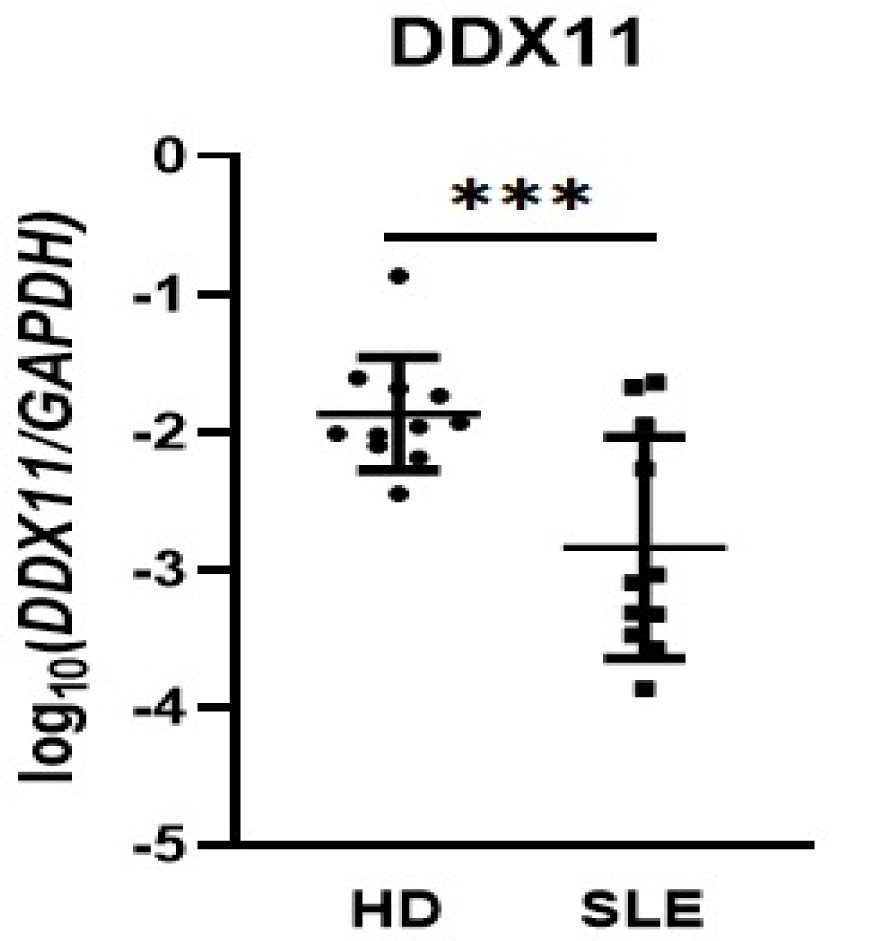
*DDX11* qPCR of monocytes from SLE and healthy donors. qPCR confirmation of *DDX11* expression in monocytes isolated from 12 SLE patients and 12 healthy donors. DDX11 is downregulated in SLE patients. Data was adjusted as logarithms. Logarithmic Normalized DDX11/GADPH: unpaired t-test, P= 0.0010. DDX11/GAPDH: Mann Whitney test, P= 0.0096.

### Protein network analysis

Protein network analysis using STRING of 187 (55+132) genes with significant differential expression in GEO datasets showed a complex interaction network. *DDX11* interacted indirectly with several genes that were found to be significant in OASIS loci and GEO datasets. *DDX11* interaction with *STAT1/STAT4* was mediated via *RIF1, RBM25* and then *BAZ1A* (Figure S1a). Interaction analysis of 55 genes having significant differential expression in at least 2 GEO datasets and with their expression directionally aligned, showed that *DDX11* directly interacted with *MED6* (interaction score 0.31) (Figure S1b). Interestingly, the remaining 132 genes included several known SLE genes such as *IRF5, BLK, TNIP1* and *CD44*, however these genes were either significant in a single GEO dataset only or did not have a uniform up-/ down-regulation for the significant probes.

## Discussion

This study employed a genetic and functional analysis using OASIS, a locus-based test, and multiple gene expression datasets to identify novel SLE genes. The most important finding of this study was the identification of *DDX11* which was significantly downregulated in both the GEO (P=3.60E-02) and the Cordoba (P=8.02E-03) SLE expression datasets. *DDX11* downregulation in SLE was confirmed (*P*=0.001) using qPCR on monocytes. The most significant SNP rs3741869 in OASIS locus 19 containing the gene *DDX11*, was found to be a *cis*-eQTL regulating the expression of *DDX11* (P=8.62E-05). *DDX11* interacted with multiple known SLE genes including *STAT1/STAT4*, identified using genomic convergence of OASIS loci and expression analysis.

DEAD/H (Asp-Glu-Ala-Asp/His) box helicase, *DDX11* is a RNA helicase thought to be involved in genome stability. It is located at the 12p11.21 locus that associated with SLE in two GWAS. Moreover, *DDX11* was found to be downregulated in SLE in multiple expression datasets and this finding was confirmed with qPCR in SLE monocytes. These findings confirm *DDX11*, a DNA helicase, as a novel SLE gene. *DDX11* mutations cause the Warsaw Breakage Syndrome (WABS), an autosomal recessive cohesinopathy with the clinical triad of severe congenital microcephaly, growth restriction, and sensorineural hearing loss (Eppley et al., 2017). WABS shows features of genome instability similar to Fanconi anemia (FA). Several genes are involved in the FA pathway which is critical to DNA repair of interstrand crosslinks (ICL). *DDX11* functions as a FA pathway backup and its deficiency leads to impaired ICL repair (Abe et al., 2018). Moreover, *DDX11* plays an important role in B-cell immunoglobulin (Ig) diversification (Abe et al., 2018) which makes it an even more interesting candidate gene for SLE. Inhibition of *DDX11* expression led to extensive apoptosis in melanoma cells (Bhattacharya et al. 2012). Previously, another melanoma associated gene, Melanoma differentiation antigen 5 (MDA5), has been shown to be important in SLE (Saeed, 2017a).

Though *DDX11* was the only gene the present study could confirm as significant in SLE, we identified several other candidate genes as well. These include *NRF1, TTC9* and *BARHL1*, which need additional data for verification. Hence, replication based genomic convergence approach with OASIS, and expression studies can help identify novel SLE genes. Measuring *DDX11* expression by qPCR in monocytes from SLE patients as described here, can potentially be used as a clinically useful predictive marker for SLE. This may allow early recognition and treatment of SLE possible and may significantly impact prevention of SLE complications.

## Methods

### Datasets

GWAS datasets were obtained online from dbGAP repository. Meta-analysis of two SLE datasets, phs000202 (Harley et al 2008) and phs000122 (Hom et al. 2008), was conducted using OASIS (Saeed, 2017b). The dataset phs000202 consisted of 706 SLE females and 353 controls and was used for screening (Harley et al 2008), while phs000122, comprising of 1435 SLE cases and 3583 controls genotyped for 500K single nucleotide polymorphisms (SNPs), was used as the replication dataset (Hom et al. 2008). SNIPPER was used to identify genes in OASIS loci. This list of reference genes was tested in 3 GEO datasets (GSE30153, GSE13887, GSE10325) for expression. Two datasets (GSE30153, GSE13887) had SLE cases and healthy controls whereas GSE10325 had three cellular fractions (T, B cells and Monocytes) resulting in 3 additional datasets (GSE10325: T, B, M). A fourth gene-expression dataset, GSE50395, from Cordoba, Spain was used for validation (Perez-Sanchez et al, 2015). However, the expression analysis comparing SLE with healthy controls is new. Figure 1 details the methodology for gene discovery.

### eQTL Analysis and qPCR

Significant SNPs in loci harboring differentially expressed genes in SLE were tested using Blood eQTL Browser (expression quantitative trait loci) (Westra et al., 2013). We confirmed the expression of *DDX11* using qPCR on monocytes of 12 healthy donors and 12 SLE patients from Cordoba, Spain. Specific primers for DDX11 were previously reported (Hormaechea-Agulla et al, 2017).

### Protein network analysis

All genes with significant and consistent directional change in expression in the GEO datasets were tested for protein–protein interactions using STRING (Szklarczyk et al., 2015). Gene pairs that were either co-expressed or involved in experimentally validated interactions with at least a low score (0.15) were evaluated.

## Conflict of Interest

None

## Acknowledgements

The authors thank all patients and healthy subjects for participation in the study. This work was accepted as a Poster in the 12^th^ European Lupus Meeting. Due to the COVID19 global outbreak the Meeting was canceled and this work could not be presented, however, the Abstract was recently published http://dx.doi.org/10.1136/lupus-2020-eurolupus.136

## Contributors

All authors were involved in drafting the article or revising it critically for important intellectual content, and all authors approved the final version to be published.

## Funding

This study was supported by grants from the Instituto de Salud Carlos III (PI18/00837), cofinanciado por el Fondo Europeo de Desarrollo Regional de la Unión Europea “Una manera de hacer Europa,” Spain, and the Spanish Inflammatory and Rheumatic Diseases Network (RIER), Instituto de Salud Carlos III (RD16/0012/0015). C.L-P was supported by a contract from the Spanish Junta de Andalucía (“Nicolas Monardes” program).

**As in Acknowledgements**

## Competing interests

None declared.

## Patient consent for publication

Not required. The study was approved by the Ethical Research Committee of Hospital Universitario Renia Sofia.

## Provenance and peer review

Not commissioned; externally peer reviewed.

## Data availability statement

All data relevant to the study are included in the article or uploaded as supplementary information. All datasets are in publically available repositories with their accession numbers in the manuscript.

## Web Resources

OASIS software (Python 2.7.9 code): https://github.com/dr-saeed/OASIS/blob/master/OASIS.py

SNIPPER: http://csg.sph.umich.edu/boehnke/snipper/

Blood eQTL Browser: https://genenetwork.nl/bloodeqtlbrowser/

STRING: http://string-db.org

## Legends

**Figure S1.**
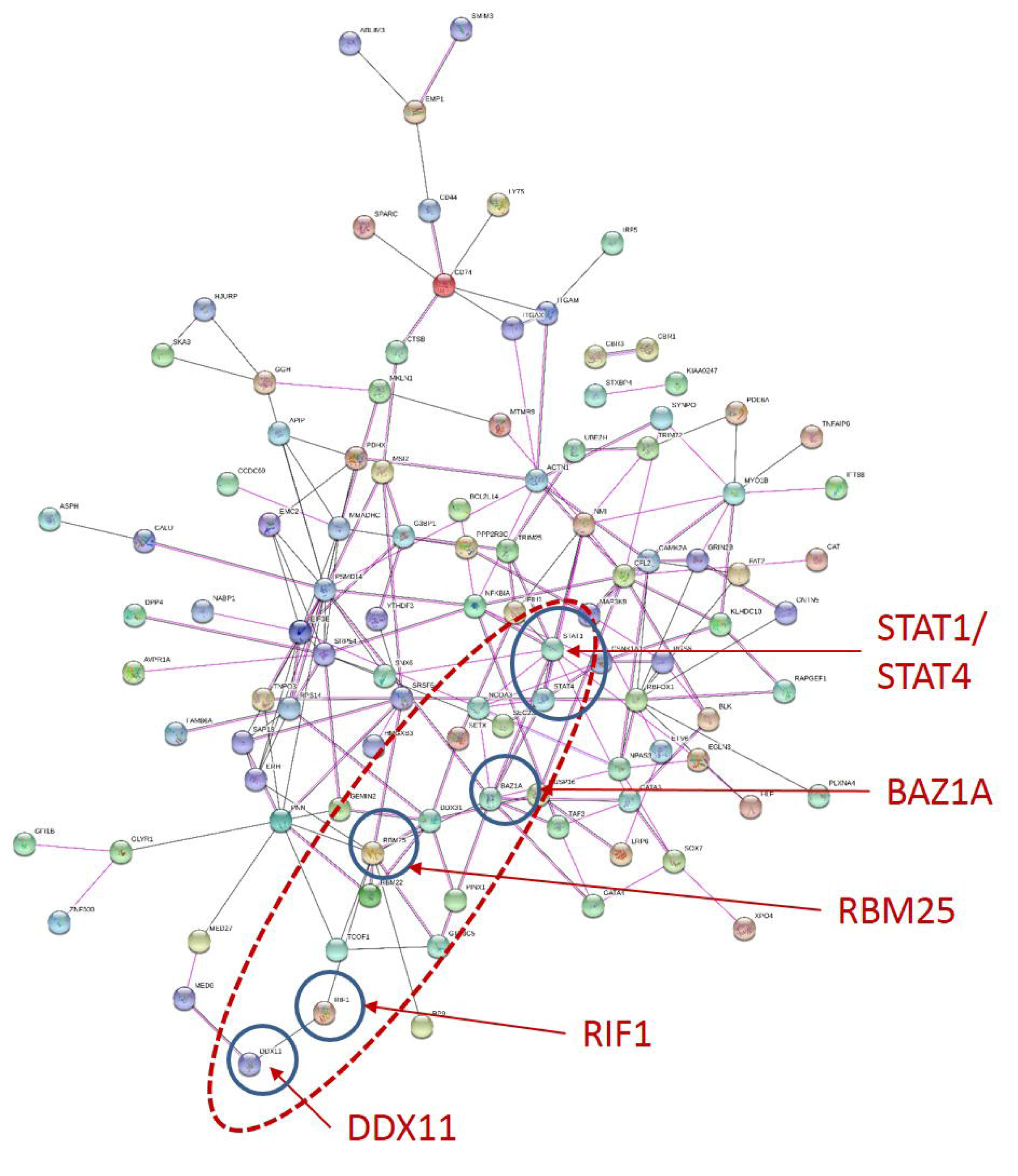
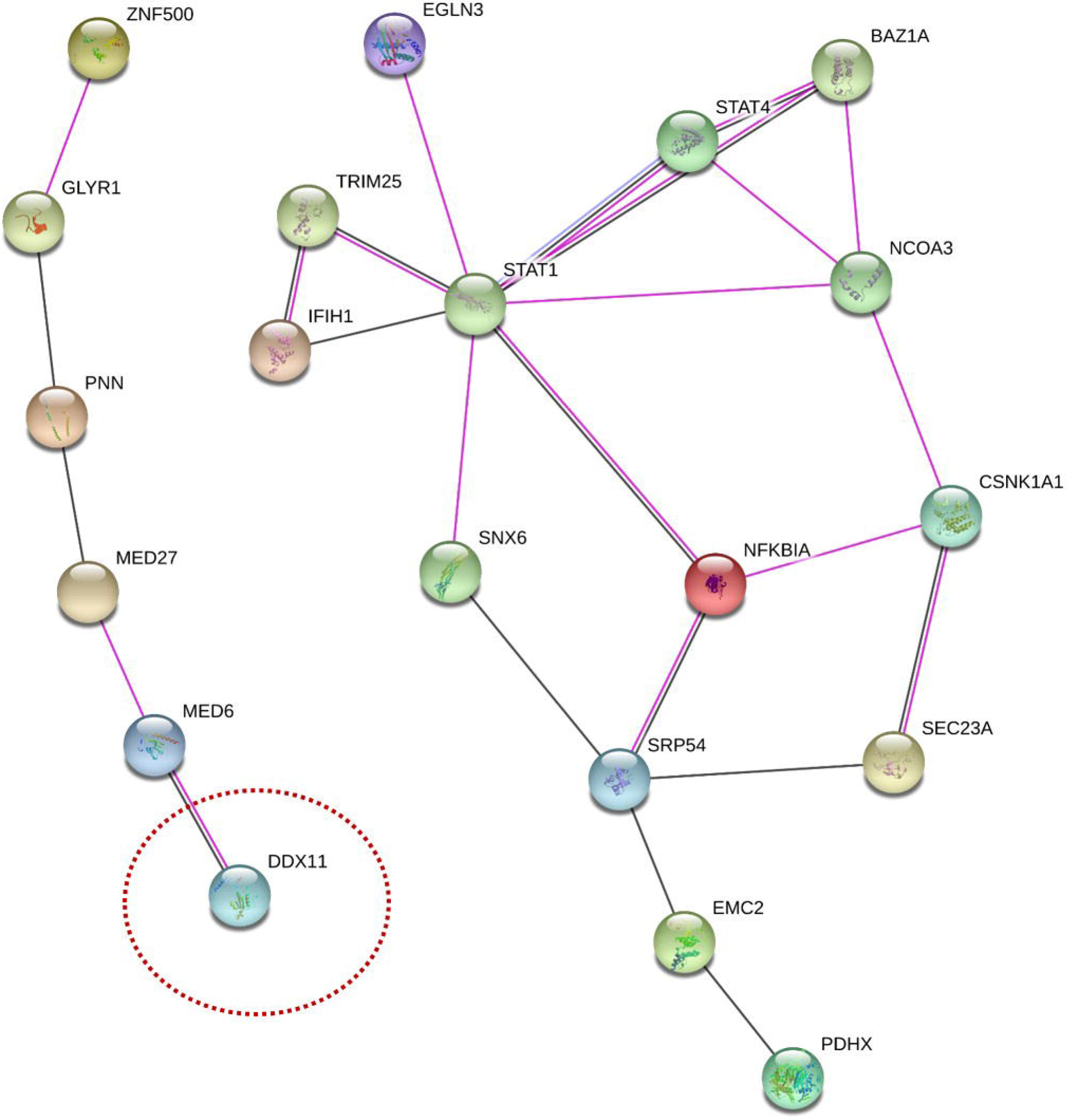
*DDX11* Protein network analysis. STRING protein network analysis of 187 unique genes (S1a) that had significant change in expression in 6 GEO datasets and were located in the 30 SLE loci. STRING analysis of 55 genes (S1b) having significant differential expression in at least 2 GEO datasets and with their expression directionally aligned. Minimum interaction score of 0.15 was chosen for either Co-expression or Experimental interaction.

**Table S1. Reference list of genes in OASIS identified SLE loci**

In 30 overlapping SLE loci identified by OASIS in two GWAS datasets, 410 genes were found using SNIPPER

**Table S2. Genomic Convergence of the reference genes in GEO Expression Datasets**

In 6 GEO datasets, 187 unique genes were identified that had significant change in expression and were located in the 30 SLE loci.

**Table S3. Genes with significant and consistent expression change in at least 2 GEO datasets**

Of the 187 candidate genes 55 were found to be differentially expressed in at least 2 GEO datasets with all probes directionally aligned.

**Table S4.** Expression quantitative trait locus (eQTL) analysis of significant genes in Cordoba validation dataset. eQTLs were found in 6 significantly expressed genes in the Cordoba dataset. DDX11 expression was modulated by the SNP rs3741869 that showed maximum significance in the DDX11 genomic locus (OASSI locus 19). The other eQTL SNPs modulated expression of genes not found to be significant in this study. Interestingly *ITGAM/ITGAX*, which are known SLE genes, were regulated by the SNPs rs11574637 and rs9888739, located in *TRIM72* and functioned as cis-eQTLs.

| OASIS Locus | Gene | cis / trans | P-value | SNP | Probe | SNP Alleles | Z-score | Modulated Gene |
| --- | --- | --- | --- | --- | --- | --- | --- | --- |
| 13 | *BARHL1* | trans | 3.21E-06 | rs11243676 | 5900274 | G>A | 4.66 | EDA |
| 13 | *BARHL1* | cis | 2.93E-06 | rs1185995 | 1340114 | G>A | 4.68 | TTF1 |
| 18 | *CDKN1B* | cis | 1.37E-03 | rs7980903 | 2060273 | T>C | -3.20 | BCL2L14 |
| 18 | *LOH12CR2* | cis | 1.37E-03 | rs7980903 | 2060273 | T>C | -3.20 | BCL2L14 |
| 19 | *DDX11* | cis | 8.62E-05 | rs3741869 | 4890750 | G>A | 3.93 | *DDX11* |
| 25 | *ADAM21* | cis | 1.10E-19 | rs7141009 | 4900164 | T>G | 9.08 | - |
| 25 | *ADAM21* | cis | 5.55E-14 | rs7141009 | 3170093 | T>G | -7.52 | KIAA0247 |
| 25 | *ADAM21* | cis | 4.31E-19 | rs2052219 | 940129 | G>A | -8.93 | WDR21A |
| 27 | *TRIM72* | trans | 1.10E-05 | rs11574637 | 4920242 | T>C | -4.40 | - |
| 27 | *TRIM72* | cis | 9.91E-33 | rs11574637 | 4490500 | T>C | 11.91 | ITGAX |
| 27 | *TRIM72* | cis | 2.18E-06 | rs11574637 | 1710070 | T>C | 4.74 | AC093520.4,ITGAM |
| 27 | *TRIM72* | cis | 1.31E-22 | rs9888739 | 4490500 | C>T | 9.79 | ITGAX |
| 27 | *TRIM72* | cis | 6.56E-06 | rs9888739 | 1710070 | C>T | 4.51 | AC093520.4,ITGAM |
| 27 | *TRIM72* | cis | 2.11E-03 | rs9888739 | 5220347 | C>T | 3.07 | BCKDK |

